# *In vivo* generation of heart and vascular system by blastocyst complementation

**DOI:** 10.1101/2022.10.04.510637

**Authors:** Giulia Coppiello, Paula Barlabé, Marta Moya-Jódar, Gloria Abizanda, Carolina Barreda, Elena Iglesias, Javier Linares, Estibaliz Arellano-Viera, Adrian Ruiz-Villalba, Eduardo Larequi, Xonia Carvajal-Vergara, Beatriz Pelacho, Felipe Prósper, Xabier L. Aranguren

**Author notes:** these authors contributed equally (Giulia Coppiello, Paula Barlabé).

## Abstract

The generation of organs from stem cells by blastocyst complementation is a promising approach to cover the clinical need for transplants. In order to generate rejection-free organs, complementation of both parenchymal and vascular cells must be achieved, as endothelial cells play a key role in graft rejection. Here we used a lineage-specific cell ablation system to produce mouse embryos unable to form both the cardiac and vascular systems. By mouse intraspecies blastocyst complementation we rescued heart and vascular development separately and in combination, obtaining complemented hearts with cardiomyocytes and endothelial cells of exogenous origin. Complemented chimeras were viable and reached adult stage, showing normal cardiac function and no signs of histopathological defects in the heart. Furthermore, we implemented the cell ablation system for rat-to-mouse blastocyst complementation, obtaining xenogeneic hearts whose cardiomyocytes were completely of rat origin. These results represent an advance in the experimentation towards the *in vivo* generation of transplantable organs.

## INTRODUCTION

Organ transplantation is the ultimate treatment for end-stage organ diseases. However, shortage of transplantable organs is a recognized major public health issue worldwide^1^. In this scenario, one of the latest ambitions of regenerative medicine is to generate human organs from pluripotent stem cells (PSCs). One promising approach that has been investigated in the last decade is the *in vivo* generation of exogenous organs in animal recipients by blastocyst complementation^2^. This technique consists in the microinjection of PSCs in mutant preimplantation embryos unable to develop a given organ, where exogenous cells can colonize the empty developmental niche and respond to the environmental cues to restore organ development^3^. For that, gene edited dysorganogenetic embryos are needed, as well as PSCs with chimeric capacity^3,4^. Proofs of concept have showed that interspecies blastocyst complementation is feasible in closely related species such as rat and mouse^4,5,6,7,8,9^ and that human PSCs hold the potential to contribute to pig development and generate post-implantation human-pig chimeras^9,10,11^. Furthermore, intraspecies blastocyst complementation of mouse models has allowed the *in vivo* generation of several organs^3^. Nonetheless, while restored organs presented parenchymal cells of complete exogenous origin, stromal and endothelial cells were a mix of donor and host cells^3,4^. Otherwise, the vascular and hematopoietic systems formation has been rescued intraspecies both in Flk1^12,13^ and Etv2^14^ knockout (KO) models. However, as endothelial cells (ECs) play a pivotal role in the acute and hyperacute rejection both in allotransplantation and xenotransplantation^15,16^, the simultaneous substitution of the parenchymal and vascular cells is needed to generate rejection-free organs^12,4^. The most widespread approaches to generate organ-disabled embryos for blastocyst complementation are based on the biallelic disruption of a master regulatory gene of a specific cell lineage by crossing heterozygous mutant mice (efficiency of KO production: 25% of the progeny). However, crossing double heterozygous mice to impair the formation of both parenchymal and endothelial cells would result in a double KO production efficiency of only 6.25%. Alternatively, microinjection of CRISPR/Cas9 system in 1-cell stage embryos has been proposed^9^. This technology gives variable efficiency of KO production but with the main drawback of uncontrolled generation of mosaic embryos, especially if several genes are disrupted. Therefore, an efficient and reliable method to produce parenchymal and endothelial cell-disabled embryos emerges as a necessity to advance in the generation of rejection-free organs.

On the other hand, the heart is an organ that has been elusive to blastocyst complementation with the KO models, as the mutation of master regulators for cardiogenesis has failed to create an empty developmental niche in the heart^3^. Heart formation relies on the interaction of different key genes and neither the disruption or deletion of Gata4^19,20^, Nkx2.5^9,21^ or Mesp1^22^, or the double KO of Id1 and Id3^23^, nor the expression of a dominant-negative form of Tbx5^24^ resulted in a heart agenesis phenotype. Accordingly, when Mesp1/ Mesp2 double KO heart development has been rescued by morula aggregation, endogenous cells were able to contribute to the atria^25^. Similarly, cardiac defects and midgestational lethality of Id1/ Id3 double KO have been rescued by blastocyst microinjection of mouse embryonic stem cells (mESCs), but the cardiac muscle composition was mosaic^23^. Finally, Wu and colleagues have rescued Nkx2.5 KO embryo development by blastocyst microinjection of rat PSCs but have obtained hearts composed by a mix of donor and host cardiomyocytes^9^.

In this study, in order to achieve the efficient generation of both cardiomyocyte and endothelial cell-deficient embryos, we employed the cell ablation system. With this model, the directed ablation of a specific cell lineage is obtained through the Cre-dependent expression of the suicide gene diphtheria toxin subunit A (DTA), where Cre expression is regulated by a chosen promoter-enhancer^26,27,28^. Here we used the Nkx2.5-Cre strain^29^ to ablate the cardiomyocyte progenitor pool and the Tie2-Cre strain^30^, to ablate the endothelial progenitor pool. The homeobox transcription factor NKX2.5 is one of the earliest markers of the cardiac lineage in vertebrates, which is expressed in the early cardiac crescent at E7.5 – 7.75 in mouse and subsequently in the secondary heart field, in cardiac progenitors contributing to the myocardium and the outflow tract^29^. When used to study the regenerative capacity of the embryonic heart, Nkx2.5-Cre x DTA - mediated cell ablation resulted in the depletion of the cardiomyocyte progenitor pool^31^. On the other hand, TIE2 is a transmembrane tyrosine kinase receptor specifically and uniformly expressed in endothelial progenitors and ECs starting from day 8 of embryonic development, as well as in some hematopoietic progenitor cells^32;33^.

Herein we combined the Nkx2.5-Cre and Tie2-Cre x DTA models to conduct intra- and interspecies blastocyst complementation studies. With mouse PSCs, we rescued simultaneously the heart and vascular system development, generating a heart with cardiac and endothelial cells of exogenous origin. Furthermore, using rat PSCs we complemented the cardiac system, forming a beating heart entirely composed by rat cardiomyocytes. These results represent an advance in the *in vivo* generation of xenogeneic hearts.

## RESULTS

### Heart complementation in Nkx2.5-Cre; R26-DTA cell ablated embryos

To corroborate the expression pattern induced by the *Nkx2.5*-IRES-Cre transgene^29^, we crossed Nkx2.5-Cre mice with Rosa26-ZsGreen (ZSG) Cre-mediated reporter mice Ai6^34^, from now on R26-ZSG. At E14.5. We detected homogeneous *Nkx2.5*-driven reporter expression in the heart and also mosaic ZSG expression in the lungs, trachea, tongue, liver and intestine (Figure S1A-B), in line with what was described before^29^. In the adult heart, virtually all cardiomyocytes were ZSG^+^ (Figure S1C), while the endothelium was mosaic, as expected, given the contribution of Nkx2.5^+^ progenitors to both cardiomyocytes and part of the heart vasculature^29,35^ (Figure S1D). Accordingly, by crossing Nkx2.5-Cre^+/+^ with R26-DTA^+/+^ ^36^ mice, we obtained Nkx2.5-Cre; R26-DTA acardiac embryos with 100% efficiency (Figure 1A and B): embryos retrieved at E9.5 lacked any heart structure and showed marked growth-retardation in comparison to age-matched controls (R26-DTA) (Figure 1B). For cardiac complementation experiments, 340 Nkx2.5-Cre; R26-DTA embryos were microinjected at E2.5 (8-cell stage) with three different GFP/ZSG labelled mouse PSC (mPSC) lines–and were transferred to 26 pseudogestant mothers. Cardiac complementation was analyzed at E14.5. At this stage, we obtained 14 embryos, all of them being complemented chimeras that were normal in size and morphology and displayed a restored beating heart (Figure 1C; Table S1). In complemented embryos, the heart was mainly derived from the ZSG^+^ exogenous cells, while the exogenous cell contribution in other tissues was mosaic (representative fluorescence image in Figure 1D). Immunostaining with cardiac troponin I (cTnI) shows that all the cardiomyocytes (CM) of complemented embryos derived from the exogenous ZSG expressing cells, while in chimeras generated from control embryos (R26-DTA), CMs had a mixed origin (Figure 1E). On the other hand, heart’s endothelial cells of complemented chimeras were derived both from donor and host embryo’s cells, as showed by the CD31 staining on coronary and capillary endothelium, the endocardium and the cardiac valves of a complemented embryo (Figure 1F). These results demonstrate that blastocyst complementation of Nkx2.5-Cre; R26-DTA embryos produces complemented hearts whose cardiomyocytes are derived entirely from donor PSCs, while endothelial cells are mosaic.

**Figure 1:**
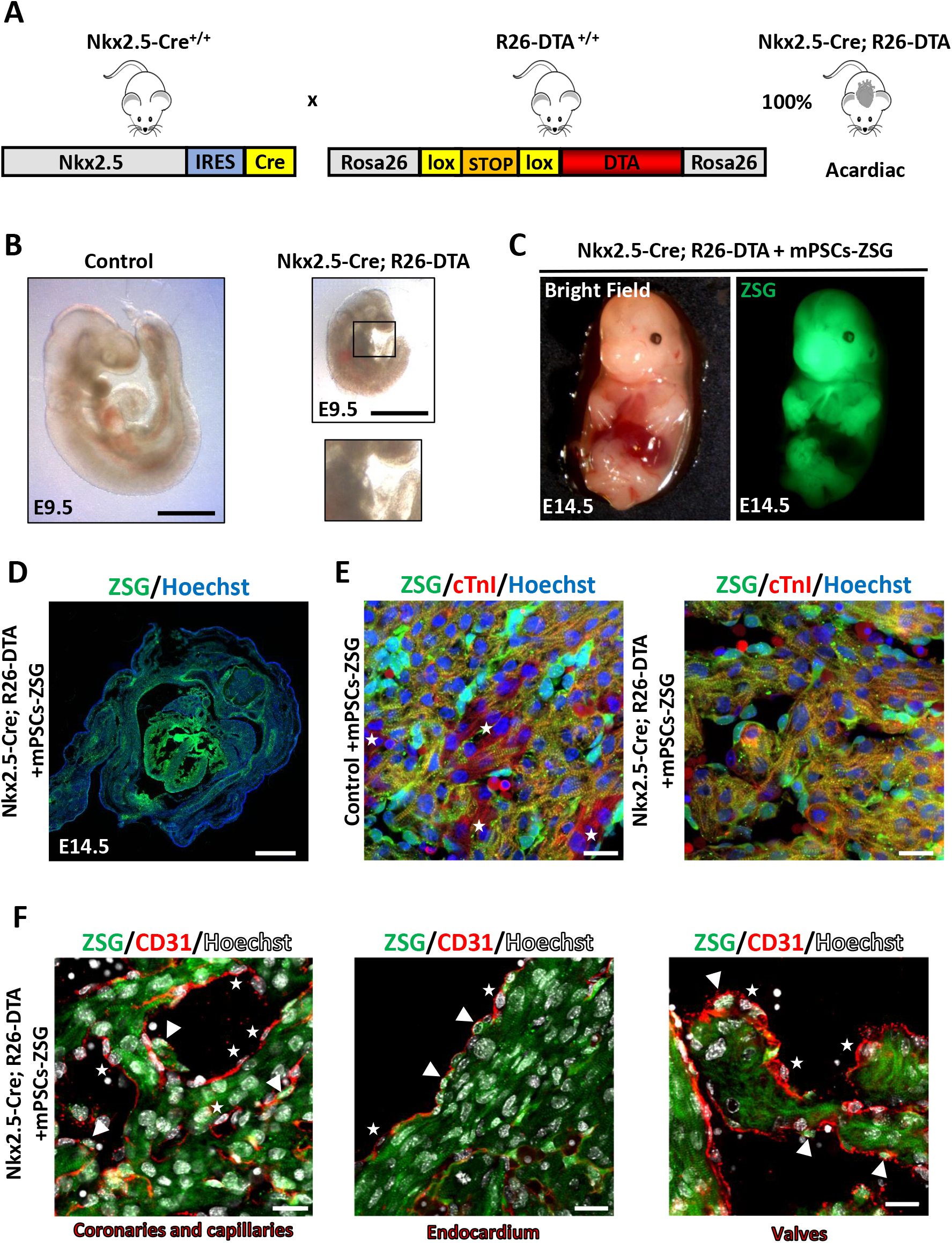
Intraspecies heart complementation. (A) Graphical representation of the transgenic mouse strains crossed to obtain acardiac embryos. (B) Whole-mount images comparing control and acardiac Nkx2.5-Cre; R26-DTA embryos at E9.5. Scale bar 1mm. Insert shows a magnification of the heart region. (C) Whole-mount bright field and fluorescence images of a complemented Nkx2.5-Cre; R26-DTA at E14.5. (D) Microscopic image of a complemented Nkx2.5-Cre; R26-DTA cross section, showing a ZSG^+^ heart. Scale bar 1mm (E) Representative cTnI (in red) immunofluorescence images of the heart of a control chimera (left) and a Nkx2.5-Cre; R26-DTA complemented chimera (right) at E14.5. Stars indicate ZSG^-^ cardiomyocytes. ZSG in green, nuclear Hoechst in blue. Scale bars 20μm. (F) CD31 endothelial staining (in red) in a Nkx2.5-Cre; R26-DTA complemented chimera’s heart at E14.5. Arrowheads indicate ZSG^+^ ECs; stars indicate ZSG^-^ ECs. Nuclear Hoechst in white. Scale bars 20μm.

### Complementation of the vascular system in Tie2-Cre; R26-DTA cell ablated embryos

To corroborate the expression pattern of the Cre recombinase, we analyzed the E14.5 embryos retrieved from the cross of Tie2-Cre^30^ mice with R26-ZSG mice. As expected^37^, virtually all ECs were positive for ZSG, as analyzed by FACS on digested E14.5 Tie2-Cre; R26-ZSG embryos (Figure S1E). We also analyzed adult mice organs by tissue immunostaining, and we found endothelial specific ZSG expression in all the organs analyzed (Figure S1F). Finally, we analyzed the whole blood of adult Tie2-Cre; R26-ZSG mice and found that around 90% of CD45^+^ hematopoietic cells were derived from *Tie2^+^* hematopoietic precursors (Figure S1G). We therefore confirmed that this *Tie2* enhancer-promoter drives Cre expression in all endothelial cells as well as in most of the hematopoietic cells. To produce complementable avascular host embryos we crossed hemizygous Tie2-Cre^+/-^ mice (homozygous mice are not viable due to the disruption of several genes induced by transgene insertion^30^) with homozygous R26-DTA^+/+^ mice, obtaining vascular agenesis in 50% of the embryos (Figure 2A and B). Analysis of the offspring at E9.5, revealed a marked growth retardation in Tie2-Cre; R26-DTA embryos compared to their R26-DTA littermates (Control) (Figure 2B). In total, we microinjected 382 embryos and vascular complementation was analyzed at E14.5. We retrieved 42 chimeras, 5 of which were Tie2-Cre; R26-DTA complemented embryos with normal size and morphology (Figure 2C; Table S1). In contrast with control chimera embryos, which showed mosaic origin of ECs by immunofluorescence staining, complemented Tie2-Cre; R26-DTA embryos showed complete exogenous origin of ECs (ZSG^+^) in all analyzed organs, both at capillary level and in major vessels (Figure 2D). Conversely, in both control and complemented chimeras, parenchymal cells had a mixed origin (Figure 2D). To confirm EC complementation, we quantified ZSG^+^ ECs by FACS analysis of digested Tie2-Cre; R26-DTA embryos, finding that virtually 100% of ECs had an exogenous origin, while in control littermates the percentage of ZSG^+^ ECs was similar to the global chimerism level of the embryos (Figure 2E). These results show that blastocyst complementation of Tie2-Cre; R26-DTA embryos produces chimeric embryos whose endothelial cells are completely derived by donor PSCs.

**Figure 2:**
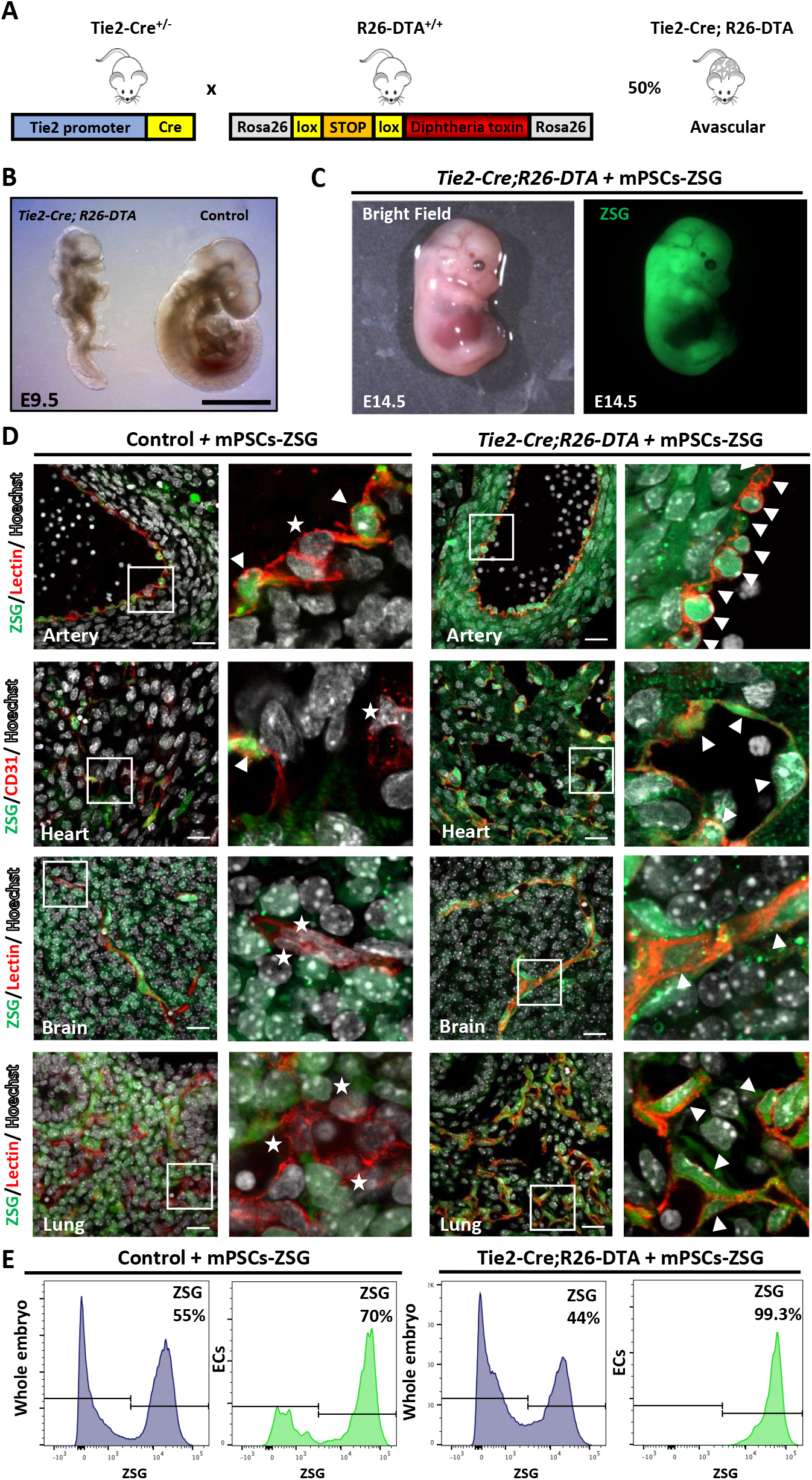
Intraspecies complementation of the vascular system. (A) Graphical representation of the transgenic mouse strains crossed to obtain avascular embryos (B) Whole-mount images comparing avascular Tie2-Cre; R26DTA and control embryos at E9.5. Scale bar 1mm. (C) Whole-mount bright field and fluorescence images of a complemented Tie2-Cre; R26DTA at E14.5 (D) Representative immunofluorescence images of an artery, the heart, brain and lung of a control chimera and a Tie2-Cre; R26DTA complemented chimera at E14.5. Endothelial BS-I Lectin or CD31 in red, ZSG in green, nuclear Hoechst in white. Arrow heads indicate ZSG^+^ ECs; stars indicate ZSG^-^ECs. Scale bars 20μm. (E) Representative histograms of the flow cytometry analysis of the chimerism in the whole embryo and in the CD31^+^; CD45^-^ ECs’ fraction from control chimeric embryos and complemented Tie2-Cre; R26DTA embryos at E14.5.

### Concomitant complementation of the heart and vascular system in Nkx2.5-Cre; Tie2-Cre; R26-DTA cell ablated embryos

To ablate simultaneously both cardiac and endothelial lineages, Nkx2.5-Cre^+/+^; Tie2-Cre^+/-^ mice were crossed with R26-DTA^+/+^ mice. This cross gives 50% of embryos that are depleted of both heart and vessels and 50% of embryos that are heart depleted only. In a first instance we assayed the two lineages complementation at the embryonic stage E14.5. From 326 microinjected embryos we retrieved 14 complemented embryos, 12 of which were Nkx2.5-Cre; R26-DTA and 2 were Nkx2.5-Cre; Tie2-Cre; R26-DTA (Table S1). To determine if complemented animals were viable after birth, we microinjected 1204 embryos and let the transferred females come to term. We obtained 21 pups with complemented heart and 11 pups with complemented heart and vascular system (Table S1). Thirteen complemented mice were grown to adulthood, showing that both heart and cardiovascular complementation supported postnatal development and adult life (up to more than 14 months). Immunofluorescence staining on adult complemented chimeras showed that cardiomyocytes were of exogenous origin in both Nkx2.5-Cre; R26-DTA and Nkx2.5-Cre; Tie2-Cre; R26-DTA mice (Figure 3A). In contrast, ECs were entirely derived from PSCs only in the case of Nkx2.5-Cre; Tie2-Cre; R26-DTA mice, both in the heart and in other tissues, while ECs were mosaic in Nkx2.5-Cre; R26-DTA complemented mice (Figure 3B). Heart function was analyzed by echocardiography in 8 complemented chimeras. All analyzed parameters (Table S2 and Figure 3C) showed no significant differences between complemented and control animals. By histopathological analysis of the heart, we analyzed whether cardiomyocytes and ECs concomitant complementation affected heart tissue composition in adult mice. In the heart of Nkx2.5-Cre; Tie2-Cre; R26-DTA complemented chimeras we did not observe signs of fibrosis, and the cardiomyocyte size and the vascular density were comparable with control mice (Figure 3D). These results evidence that it is possible to complement simultaneously the heart and vascular system in Nkx2.5-Cre; Tie2-Cre; R26-DTA embryos. Complemented chimeras grow to adulthood and present normal cardiac tissue composition and function, and their restored heart is composed by cardiomyocytes and endothelial cells entirely derived from exogenous PSCs.

**Figure 3:**
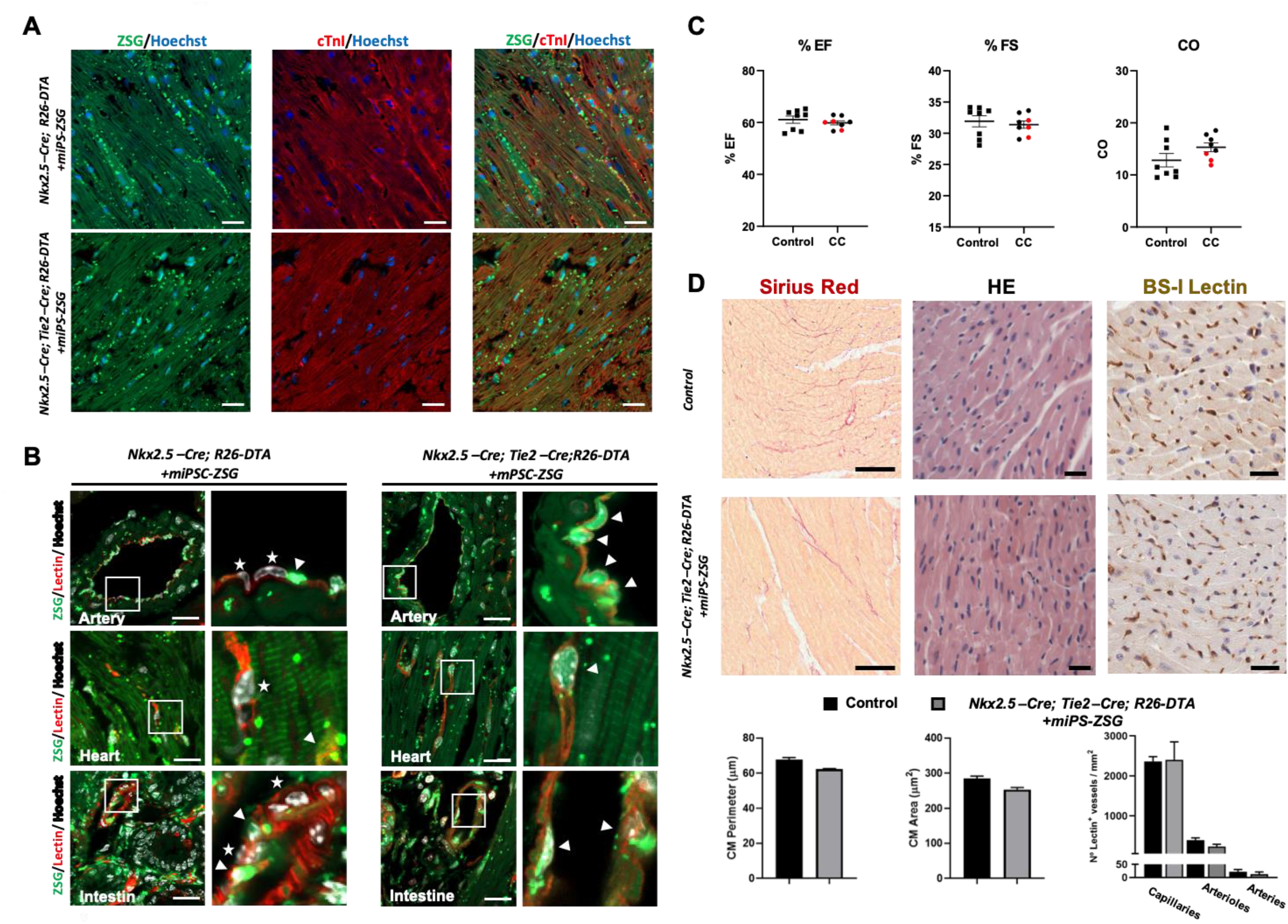
Concomitant complementation of the heart and vascular system with mPSCs. (A) Representative immunofluorescence pictures of heart sections from Nkx2.5-Cre; R26-DTA complemented chimera (on top) and from Nkx2.5-Cre; Tie2-Cre; R26-DTA complemented chimera (below) stained with cTnI (in red). ZSG in green, nuclear Hoechst (blue). Scale bars 20μm. (B) BS-I Lectin immunostaining (red) of arterial ECs, cardiac ECs and intestinal ECs of Nkx2.5-Cre; R26-DTA complemented mice (left) and of Nkx2.5-Cre; Tie2-Cre; R26-DTA complemented mice (right). Nuclear Hoechst (white). Scale bar 20μm. Arrowheads indicate ZSG^+^ ECs; stars indicate ZSG^-^ ECs. (C) Dot plot graph of ejection fraction (%EF), fractional shortening (%FS) and cardiac output (CO; ml/min) echocardiographic parameters. Dots represent complemented chimera (CC) mice: Nkx2.5-Cre; Tie2-Cre; R26-DTA mice are evidenced in red. Black squares represent control mice (R26-ZSG^+^). (n=8 per group). P value= ns. (D) Representative images of control (on top) and Nkx2.5-Cre; Tie2-Cre; R26-DTA mice hearts (below) stained for Sirius red, scale bar 50μm; H-E and BS-I Lectin, scale bar 25μm. Bar graph representation of the CM area, CM perimeter and vascular density across different vascular beds in the heart of both control (n=4) and complemented Nkx2.5-Cre; Tie2-Cre; R26-DTA (n=3). P value= ns.

### Interspecies rat-to-mouse heart complementation in Nkx2.5-Cre; R26-DTA cell ablated embryos

Our next goal was to complement the heart and vascular system in an interspecies setting using rat PSCs (rPSCs). For that, rat ESCs from eGFP Sprague-Dawley (SD) rats were injected into 278 mouse 8-cell stage embryos resulting from the cross of Nkx2.5-Cre^+/+^; Tie2-Cre^+/-^ and R26-DTA^+/+^ mice (Table S3). Microinjected embryos were transferred into the uteri of 28 mice and 14 rat-to-mouse chimeras were collected at E10.5. Among them, 4 embryos presented fully complemented looped hearts (Figure 4A; Table S3) while the other 10 chimeras had a very primitive heart structure, or were acardiac, sign of partial or absent complementation (Figure 4B). All 14 chimeras were Nkx2.5-Cre; R26-DTA and thus, only the heart, but not both heart and vascular system, was complemented in an interspecies setting, as analysed at E10.5. At E14.5 or E11.5 on the other hand, rescued embryos were not found (Table S3). A representative fluorescence microscopy image of a whole-mount Nkx2.5-stained complemented embryo shows that virtually all cardiomyocytes were rat PSC-derived, as they expressed the eGFP transgene (Figure 4C). This was validated by immunofluorescence staining on tissue sections (Figure 4D) and subsequent quantification: 99,11% of Nkx2.5^+^ cells expressed eGFP (2.010 cells analyzed). These results indicate that acardiac Nkx2.5-Cre; R26-DTA embryos can be complemented interspecies and the resulting rat-to-mouse chimeras present a restored heart whose cardiomyocytes are entirely of rat origin.

**Figure 4:**
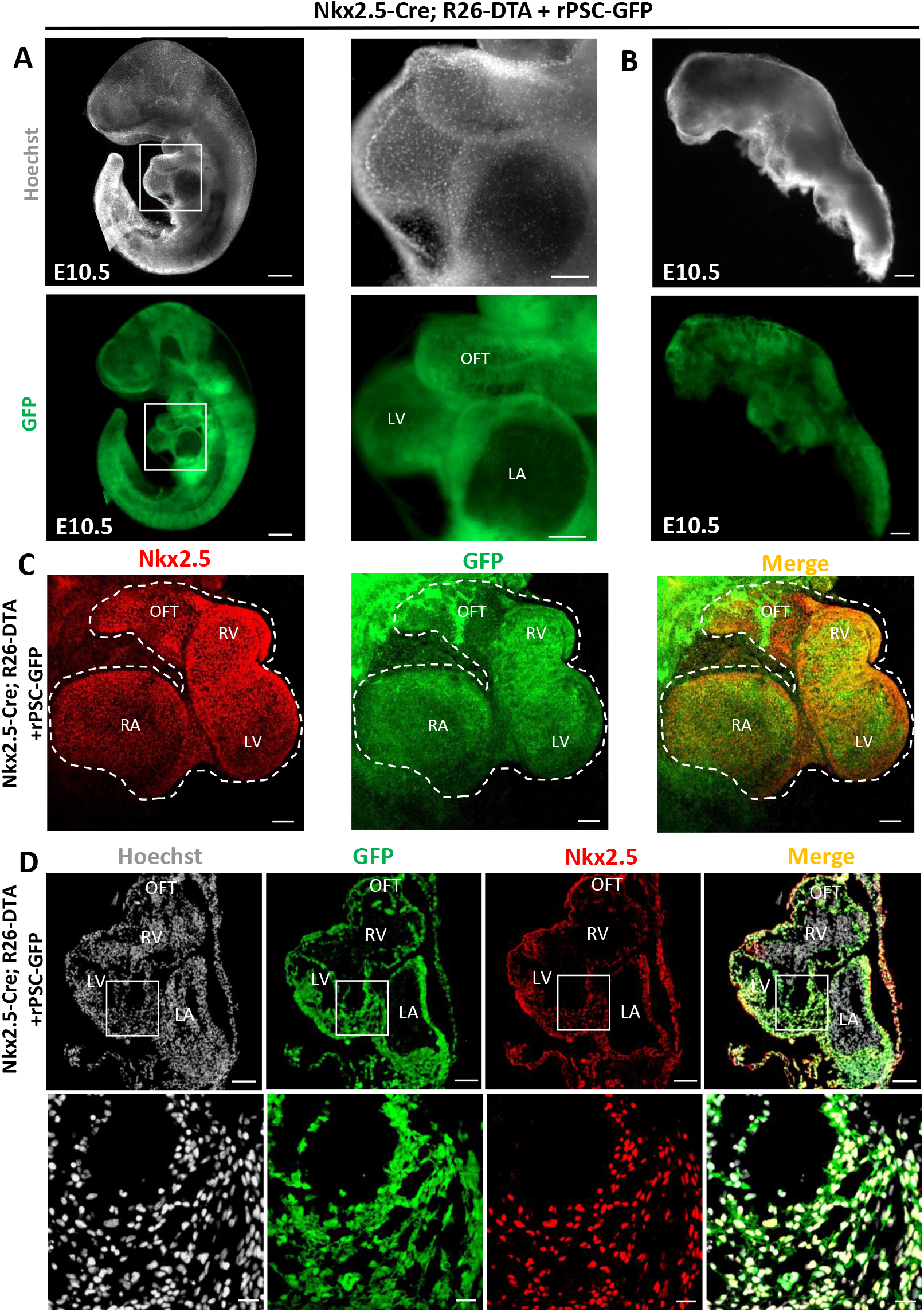
Interspecies rat-to-mouse heart complementation. Whole-mount images of Nkx2.5-Cre; R26-DTA rat-to-mouse chimeras with a complemented heart (A) and without heart complementation (B) at E10.5. Hoechst staining (white, upper panels) and GFP fluorescence (green, bottom panels). Scale bar: A, 500μm; B, 200μm. Boxed areas are enlarged in central panels. Scale bar 200μm. (C) Maximal projection of confocal stacks from the heart of a complemented Nkx2.5-Cre; R26-DTA rat-to-mouse chimera whole-mount stained for Nkx2.5 at E10.5 (in red). Dotted lines indicate heart. Scale bar, 100μm. (D) Representative immunofluorescence images of a complemented heart from a Nkx2.5-Cre; R26-DTA rat-to- mouse chimera at E10.5: Nkx2.5 (red), GFP (green) and Hoechst (white). Scale bar: 100μm. Magnified views of boxed areas are shown in bottom panels. Scale bar: 20μm. RV: right ventricle; LV: left ventricle; RA: right atria; LA: left atria; OFT: outflow tract.

## DISCUSSION

In this study, we successfully achieved concomitant blastocyst complementation of heart and vascular system, obtaining for the first time hearts whose cardiomyocytes and endothelial cells were entirely derived from exogenous mPSCs. Complemented mice reached adulthood, their hearts were normal in tissue composition and were fully functional. The concomitant generation of parenchyma and ECs *in vivo* from PSCs represents a significant advance towards the generation of rejection-free hearts for clinical transplantation, as ECs’ immunogenic epitopes and mismatched MHC-I are readily detected as foreigners and attacked by the host immune system upon (xeno)transplantation^15,16^. A similar venture was undertaken by Matsunari et al., who generated a combined pancreas–endothelium-disabled (*PDX1* and *KDR* KO) pig model and complemented it intraspecies^17^. In this case, mutated embryos were obtained by somatic cell nuclear transfer (SCNT) after CRISPR/Cas9 gene editing of fibroblasts *in vitro*. However, the SCNT approach has a limited use for the generation of organs *in vivo*, as it produces only 1-2% of viable embryos^18^. To obtain organ-disabled embryos we employed the conditional cell ablation system. This method allowed us to work with 50% to 100% of dysorganogenetic embryos. This represents a significant improvement in efficiency compared to crossing heterozygous mice, which is the standard method for producing organ-disabled embryos and gives 25% of complementable progeny when targeting a single gene and a 6.25% when targeting two genes. Noteworthily, the cell ablation system has the potential to eliminate multiple cell types with up to 100% efficiency, as there is no need for endogenous gene disruption. The implementation of this system could solve the problem of efficiently targeting all the cell types that constitute a given organ. With such models, generation of fully exogenous organs could be attempted by blastocyst complementation.

Moreover, the cell ablation system allows to directly eliminate a given progenitor cell pool, leaving a free developmental niche for exogenous PSCs to colonize. This is not the case for deletion or mutation of studied master regulators of heart development, as the phenotype of these models is not acardiac^4;19–25^. Accordingly, several chimera experiments performed using cardiac genes KO embryos have shown mixed host and donor contribution to cardiomyocytes^9;23;25^. Here, through interspecies blastocyst complementation of Nkx2.5-Cre x R26-DTA embryos we have generated hearts fully composed by rat cardiomyocytes. This represents an advance compared with previous results from Wu et al., where rPSCs rescued heart development in Nkx2.5-disrupted mouse embryos with mixed donor-host cell contribution to cardiomyocytes^9^. However, in line with this study^9^, in interspecies setting we observed the loss of Nkx2.5-Cre; R26-DTA complemented embryos beyond E10.5. Another drawback that we experimented in interspecies blastocyst complementation experiments was the failure to simultaneously complement the heart and vascular system. This might be due to the lack of interspecies vascular complementation per se, as others have reported that vascularization by rat cells was only partially achieved until E9.5^12^ and E10.5^13^ in Flk1 KO models. Both the loss of embryos with complemented hearts beyond E10.5 and the missed complementation of heart and vascular system at the same time might be due to interspecies barriers that lead to developmental incompatibilities. Although a comprehensive understanding of these barriers is still lacking, different mechanisms have been proposed: inappropriate timing and site for cellular interactions, mismatched ligand-receptor interactions, affinity differences in adhesion molecules, and cell-competition induced cell death^38;39;40^. Exploring all these processes could clarify the nature of these barriers, the timing of their action and the tissues involved, as interspecies adult mouse chimeras with functional rat pancreas^5^ and thymus^6^ have been successfully generated, and therefore it appears to be a tissue-dependent process. In summary, in this study, we generated complemented hearts with both cardiomyocytes and endothelial cells of exogenous origin by blastocyst complementation of cell ablated embryos, providing an advance towards rejection-free *in vivo* organ generation. Our data also evidence that interspecies heart complementation is achievable at embryonic level, paving the way for xenogeneic heart generation.

## ACKNOWLEDGMENTS

We thank Francisco José Rodríguez Díaz, Belen Pintado and Verónica Domínguez Plaza for technical advice on embryo manipulation. Dr. José María Pérez-Pomares for his help with the morphological characterization of rat-to-mouse complemented heart. Natalia Aguado for technical support. Dr. Irantzu Llarena for the assistance with confocal imaging. CIMA Morphology and Imaging units for the assistance provided with histological samples preparation and images recording and with echocardiography recording and analysis. CIMA Flow cytometry unit for the assistance provided with flow cytometry and cell sorting.

This work was supported by RETIC TerCel (RD16/0011/0005), Marie-Curie Fellowship (FP7-PEOPLE-2013-IEF 626220), the Ramón y Cajal State Program (RYC-2015-17233, financed by MCIN/AEI and FSE), Proyecto de I+D Retos Investigación (UE/RTI2018-064485-B-I00, financed by MCIN/AEI and FEDER), Juan de la Cierva Formación Fellowship (FJCI-2014-22909, financed by MCIN/AEI and FEDER); Juan de la Cierva Incorporación Fellowship (IJCI-2017-33070, financed by MCIN/AEI and FEDER); FIMA predoctoral fellowship.

## AUTHOR CONTRIBUTION

G.C conceived the research, provided funding, performed mouse-to-mouse experiments, analyzed the data, and wrote the manuscript. P.B. conceived the research, provided funding, performed rat-to-mouse experiments, analyzed the data, and wrote the manuscript. G.A. performed embryo transfer and vasectomization, and edited the manuscript; M.M, C.B, E.L. and E.I. provided assistance with experiments and edited the manuscript. J.L. and E.A. generated and characterized miPS cell lines and edited the manuscript. A.R.V provided assistance with imaging and edited the manuscript, X.C.V provided materials and technical advice and edited the manuscript; B.P. collected and analyzed histological data and edited the manuscript; F.P. provided funding and edited the manuscript. X.A. conceived the research, provided funding supervised the experiments, analyzed the data, and wrote the manuscript.

## DECLARATION OF INTERESTS

The authors declare no competing interests.

## STAR METHODS SECTION

### RESOURCE AVAILABILITY

#### Lead Contact

Further information and requests for resources and reagents should be directed to and will be fulfilled by the Lead Contact, Xabier L. Aranguren.

#### Material availability

Mouse and rat PSCs generated in this study are available from the Lead Contact’s laboratory upon request and following the completion of a Material Transfer Agreement.

#### Data and Code Availability

This study did not generate any code or dataset.

### EXPERIMENTAL MODEL AND SUBJECT DETAILS

#### Animals

Nkx2.5-Cre mice (*B6.129S1(SJL)-Nkx2-5^tm2(cre)Rph^*/J; stock n°024637 from Jackson laboratories)^29^ and Tie2-Cre mice (B6.Cg-Tg(Tek-cre)12Flv/J, stock n° 004128, Jackson laboratories)^30^, were crossed with DBA/2JRj mice to achieve a 50% mixed background. Homozygous R26-ZSG (Ai6; *B6.Cg-Gt(ROSA)26Sor^tm6(CAG-ZsGreen1)Hze^/J;*stock n° 007906) females were crossed with Nkx2.5-Cre^+/+^ and Tie2-Cre^+/-^ male mice to generate respectively Nkx2.5-Cre;R26-ZSG and Tie2-Cre;R26-ZSG reporter embryos and mice. Homozygous R26-ZSG were crossed with Sox2-Cre (*B6.Cg-Edil3^Tg(Sox2-cre)1Amc^*/J; stock n°008454) hemizygous mice to remove the floxed stop codon in all embryo’s cells in R26-ZSG^+/-^; Sox2-Cre^+/-^ and then backcrossed with R26-ZSG^+/-^ to select Sox2-Cre^-^ mice constitutively expressing ZSG (R26-ZSG^+^) as control mice for echocardiography and histopathological analysis. R26-DTA mice (B6;129-*Gt(ROSA)26Sor ^tm1(DTA)Mrc^*/J; stock n°010527, also purchased from Jackson laboratories) were crossed with DBA/2JRj mice (Janvier) to achieve a 50% mixed background.

Homozygous R26-DTA^+/+^ females were crossed with Nkx2.5-Cre^+/+^ male mice to produce acardiac embryos for blastocyst complementation experiments at a ratio of 100%. Hemizygous Tie2-Cre^+/-^ male mice were crossed with R26-DTA^+/+^ females to produce avascular embryos for blastocyst complementation experiments at a ratio of 50%. Nkx2.5-Cre^+/+^ mice were crossed with Tie2-Cre^+/-^ mice for two generations to obtain Nkx2.5-Cre^+/+^; Tie2-Cre^+/-^ stud males, that were crossed with R26-DTA^+/+^females to obtain 50% acardiac and 50% acardiac and avascular embryos for blastocyst complementation experiments. Embryos were collected at E2.5 and microinjected with mouse or rat PSCs for blastocyst complementation experiments. Male CD1 mice (purchased from Janvier) were vasectomized at 8 weeks of age and used as sterile studs to induce pseudopregnancy in CD1 females used as microinjected embryos recipients in blastocyst complementation experiments. All animals were housed in specific pathogen free conditions with free access to food and water. All procedures were approved by the ethical committee of University of Navarra and governmental guidelines for laboratory animals care and safety were strictly followed.

#### PSCs culture and manipulation

Mouse R1 ESCs^41^ were a kind gift by Dr. Nagy’s lab. They were transduced with a lentiviral vector containing the GFP reporter (pLenti CAG IRES GFP 69047 Addgene), the GFP^+^ fraction was FACS sorted, re-plated and expanded on irradiated mouse embryonic fibroblasts (iMEFs) as described in^41^. The AHFiPS 7 mouse iPS cell line used in this study was previously characterized in^42^. The AHFiPS 9 iPS cell line was generated and characterized at CIMA, University of Navarra as described in ^42^. Both iPS cell lines were derived from male tail tip fibroblasts of Mef2c-AHF-Cre^43^ x R26-ZSG (Ai6; *B6.Cg-Gt(ROSA)26Sor^tm6(CAG-ZsGreen1)Hze^/J;* stock n° 007906). iPS cells were expanded in KSR-LIF medium: KO-DMEM (Gibco 10829-018), 15% KSR (Gibco 10828-028), Antibiotic/Antimycotic (Gibco 15240-062; 100 U/ml); Glutamax (Gibco 35050038; 2mM), NEAA (Gibco 11140-035; 1x) 2-mercaptoethanol (Gibco 31350-010; 0,1μM), 10^3^ U/ml mouse LIF (ESG1106 Sigma-Aldrich) on iMEFs. For constitutive ZSG transgene expression, cells were electroporated with the Cre recombinase-expressing plasmid pBS185 CRE (Addgene 11916). ZSG^+^ cells were FACS sorted and expanded. For blastocyst complementation experiments, CHIR99021 (Axon1386; Axon Medchem) and PD0325901 (Axon1408; Axon Medchem) were added to the cultured media at 3μM and 1 μM concentration respectively two days before microinjection. AHFiPS 7 were used at passage 10-18, AHFiPS 9 at passage 17-20 and R1 cells were used at passage 13-19. ATCrSD78 derivation and characterization was described in^44^. Other rat ESCs were generated from rat blastocysts (E4.5) obtained by crossing Sprague-Dawley (SD) eGFP rats (SD-Tg(GFP)1BalRrrc; RRRC) as described in^44^ with minor modifications. Briefly, blastocysts outgrowths were partially digested and replated and ESCs colonies showing homogeneous GFP expression were selected and expanded. Rat ESCs were maintained on iMEFs in N2B27 3i medium containing rat LIF 10^3^U/ml (Millipore), CHIR99021 1μM PD0325901 1μM and A83 inhibitor1μM. The medium was changed every two days and cells were passaged with Accutase (Gibco) every 3-5 days. Cells between passage 6 and 13 were used for blastocyst complementation experiments. For microinjection experiments, mouse and rat PSCs’ single cell suspension was obtained by Tryple (Gibco) treatment, cells were pelleted and resuspended in culture medium with Hepes buffer (20mM) and stored at 4°C until used.

### METHOD DETAILS

#### Analysis of Nkx2.5-Cre and Tie2-Cre mouse lines

##### Reporter mice

Nkx2.5-Cre^+/+^ and Tie2-Cre^+/-^ male mice were crossed with R26-ZSG^+/+^females. Next day, vaginal plug was confirmed (E0.5). At E14.5 females were sacrificed by cervical dislocation and embryos were retrieved for their analysis by FACS or by immunofluorescence. Tie2-Cre; R26-ZSG adult mice blood was retrieved by peri-orbital bleeding with a glass capillary tube and collected in an Eppendorf tube with heparin for subsequent processing and FACS analysis. Subsequently, mice were sacrificed by cervical dislocation and their organs retrieved for immunofluorescence analysis.

##### Cell ablation models

Nkx2.5-Cre^+/+^ and Tie2-Cre^+/-^ male mice were crossed with R26-DTA^+/+^ females. Next day, vaginal plug was confirmed (E0.5). At E9.5 females were sacrificed by cervical dislocation and embryos were retrieved for their morphological analysis and were subsequently genotyped.

#### Blastocyst complementation

8 to 10-week-old R26-DTA^+/+^ females were superovulated with intraperitoneal injections of 5U of PMSG and hCG with a 47h interval and immediately mated with the selected males for the different experiments. Embryos were collected at 8 cells stage (E2.5) and microinjected with 3 to 6 mPSCs or 6 rPSCs with Eppendorf TransferMan NK2 micromanipulator coupled to a Leica DMI 3000B microscope. When microinjected with mouse PSCs morulae were cultured overnight in KSOM (LGGG-050 Global supplemented with 4mg/ml BSA (A9576 Sigma Aldrich) drops overlaid with Nidoil (VNI0049; Nidacom) at 37° in 5%O_2_, 5%CO_2_ atmosphere and the morning after 15 to 30 embryos were transferred to the uterus or infundibulum of each pseudopregnant CD1 females (E2.5 or E0.5, respectively). When microinjected with rat PSCs, morulae were cultured overnight in KSOM supplemented with rat N2B27 3i medium (1:1) prior to embryo transfer. Embryos were retrieved at E10.5, E11.5 or E14.5 for analysis, or were let to develop to term to analyze the complemented mice at early (newborn) or adult stage (6 weeks). In that case, when possible, wild type embryos (CD1 or B6;DBA2 background) were co-transferred to ensure the production of a litter big enough to avoid cannibalism. Transferred females expected to give birth to complemented chimeras were fed with a diet supplemented with 9% fat content (Teklad global Envigo).

#### FACS analysis

FACS analysis was performed on single cell suspension derived from enzymatic digestion of E14.5 Tie2-Cre; R26-DTA complemented embryos and E14.5 Tie2-Cre; R26-ZSG reporter embryos, and on whole blood of adult Tie2-Cre; R26-ZSG mice upon erythrocytes lysis with 20min incubation in ammonium chloride potassium solution (ACK) lysis buffer (150mM NH4Cl, 10mM KHCO3 and 0,1mM Na2EDTA [pH 7.2-7.4]). For embryo digestion, briefly, tissues were finely minced with a scalpel in PBS containing 1,5mg/ml Collagenase I (Gibco) and then incubated at 37° for 45min. Samples were filtered through a 40μm mesh, washed with PBS and resuspended in PBS 1%BSA (A3912 Sigma-Aldrich) for staining. Antibodies used are listed in Table S4. CD31 and CD45 were used to identify endothelial (CD31^+^ CD45^-^) and hematopoietic cells (CD31^-/low^ CD45^+^). Fluorochrome labelled antibodies were incubated with the sample for 15min at room temperature. After a wash in PBS, cells were resuspended in FACS buffer (PBS 1mM EDTA, 25 mM HEPES, 1% BSA, pH7). 7AAD was used to exclude died cells. Cell staining was analyzed with BD FACSCanto™ II with BD FACSDiva software; ZSG and GFP were analyzed in the FITC channel. Sorting was performed with FACS AreaI and AreaII. Flow cytometry data were analyzed using FlowJo software.

#### Genotyping

Genotyping of embryo donors was performed on an ear plug by DNA extraction with KAPA enzyme and PCR reaction with Kapa master mix (KK7352 Kapa biosystems).

Genotyping of complemented embryos was performed on sorted GFP^-^ fraction or placental tissues by DNA extraction with KAPA enzyme and PCR reaction with Kapa master mix (KK7352 Kapa biosystems). For adult complemented chimeras, an ear plug and/or organs’ samples were used for genotyping after DNA extraction with Kapa buffer or column (A174095250 MACHEREY-NAGEL). PCR reaction was performed with Kapa master mix or by semi-nested qPCR with SYBR Green (A25742 Applied biosystems). Primers are listed in Table S5.

#### Histology and immunohistochemistry

Embryos were fixed in 4% PFA solution in PBS overnight (E14.5) or 3h (E.10.5) at 4°C. Adult mice were anesthetized with Ketamine/Xylazine mix, and shortly perfused with PBS trough a catheter inserted in the heart apex, successively, 100μl of 30mM KCl solution was injected in the right atrium to induce diastole cardiac arrest. PBS was perfused 5 min more and then the organs were dissected and fixed overnight in 4% PFA solution at 4°C. For cryopreservation, samples were dehydrated in 30% sucrose solution in PBS overnight at 4°C and then included in OCT medium (Tissue-Tek, 4583 Sakura) and stored at −80°. Tissues were sliced with a microtome at 7μm thickness. When immunostaining, each section was incubated with a blocking solution containing 1% BSA and 0,3% Triton X-100 for 1 hour at room temperature and subsequently with the primary antibody overnight at 4° and with the secondary antibody for 2 hours, at room temperature (Table S4). Images were recorded with Confocal Scanning Laser Microscope (Zeiss LSM 800) and ZEN 2.3 software or Zeiss Axio Imager M1Esc microscope equipped with ZEN Pro software. Fiji software was used to create composite images. For paraffine embedding, fixed samples were dehydrated in 70% ethanol solution in MilliQ water. Tissue blocks were sectioned with a microtome at 6μm thickness. Paraffin-embedded sections were deparaffinized with xylene and hydrated with graded ethanol. H&E staining, Sirius red staining or BSI-Lectin immunostaining were performed as described elsewhere. Antibodies used are listed in Table S4. Vascular density was quantified on 2000-2500 vessels and cardiomyocyte cross-sectional area was quantified on 100 cells by a blinded investigator, using Leica Aperio CS2 scanner and Image J software.

#### Whole-mount immunofluorescence

Immunostaining was performed as described in^45^ with minor modifications. Embryos were fixed with PFA 4% 3h at 4°C, permeabilized with 0.5% Triton X-100 30min at RT and blocked with 3% donkey serum (Sigma, D9663), 1% BSA and 0.1% Triton X-100 overnight at 4°C. Each sample was incubated with primary antibodies 48h and secondary antibodies overnight, both at 4°C (Table S4). Whole embryos were imaged with a Confocal Scanning Laser Microscope (Zeiss LSM 800) and ZEN 2.3 system software. Image J software was used to create composite images.

#### Echocardiography

Eight animals per experimental group were analyzed at 6 weeks of age. Echocardiographic images were acquired using the Vevo3100 high resolution *in vivo* ultrasound system (FUJIFILM, VisualSonics, Toronto, Canada) equipped with a real-time scanning head (MX550D) operating at a frame rate above 200 frames per second (fps). The selected transducer operates at a center frequency of 40 MHz, a focal length of 12.7 mm, and an axial resolution of 40 μm. The images were quantified using VevoLab software (v5.1.1) and, in particular, the cardiac measurement package. Briefly, mice were anesthetized with 4% isoflurane in 80% oxygen (Isoflo®, ABBOTT, Spain) and then placed on a heating platform in a supine position. Body temperature, electrocardiography (ECG), and respiratory rate were monitored by probes connected to the extremities of the animals. Subsequently, mice were depilated with depilatory cream (Veet, Reckitt Benckise, Spain) and prewarmed ultrasound gel (Quick Eco-Gel, Lessa, Spain) was applied to the chest for observation. During the examination, the isoflurane concentration was lowered to 1% to obtain constant cardiac parameters. The functional analysis of the heart was evaluated using the systolic and diastolic functions of the left ventricle (LV): LV systolic function was assessed using ejection (EF%) and shortening (SF%) fractions obtained from LV diameters measured in diastole and systole (LVID d, LVID s) of the M-mode. To obtain the classic LV M-mode window, the caliper was placed parallel to both papillary muscles in the parasternal short-axis view (PSAX). Likewise, cardiac output (CO) (ml/min), stroke volume (SV) (μl), LV mass (mg), as well as the left ventricular anterior wall (LVAW) and the left ventricular posterior wall (LVPW) thickness (mm) were quantified using this mode. LV diastolic function was assessed by pulsed-wave Doppler detection of transmitral inflow obtained from the apical LV in a four-chamber view. During the examination, the caliper was kept as parallel as possible to the mitral blood flow. E wave (mm/s), A wave (mm/s), E/A ratio, deceleration time (MVDT) (ms), ejection time from opening to closing of the mitral valve (ET) (ms), and isovolumic LV relaxation (IVRT) and contraction (IVCT) times (ms) were measured from the pulsed Doppler plots. In addition, the TEI index was obtained by applying the following expression: (IVCT+IVRT)/ET. Finally, an estimation of diastolic function was also obtained using pulsed tissue Doppler by placing the caliper on the basal portion of the mitral annulus. Negative E’ wave (mm/s), negative A’ wave (mm/s), and the E/E’ ratio were quantified from the pulsed tissue Doppler plots.

#### Quantification and statistical analysis

Data are expressed as mean ± standard error of the mean (SEM) from pooled data of 3 to 8 independent samples. For echocardiography analysis (n=8), data normality was determined by the Shapiro-Wilk and Kolmogorov-Smirnov tests and 2-tailed (un)paired Student’s *t* test or U of Mann-Whitney test was applied accordingly. For the histological analysis of the complemented chimera (n=3-4), U of Mann-Whitney test was applied. In all cases significance level was set at α<0.05.

## SUPPLEMENTAL INFORMATION

**Figure S1:**
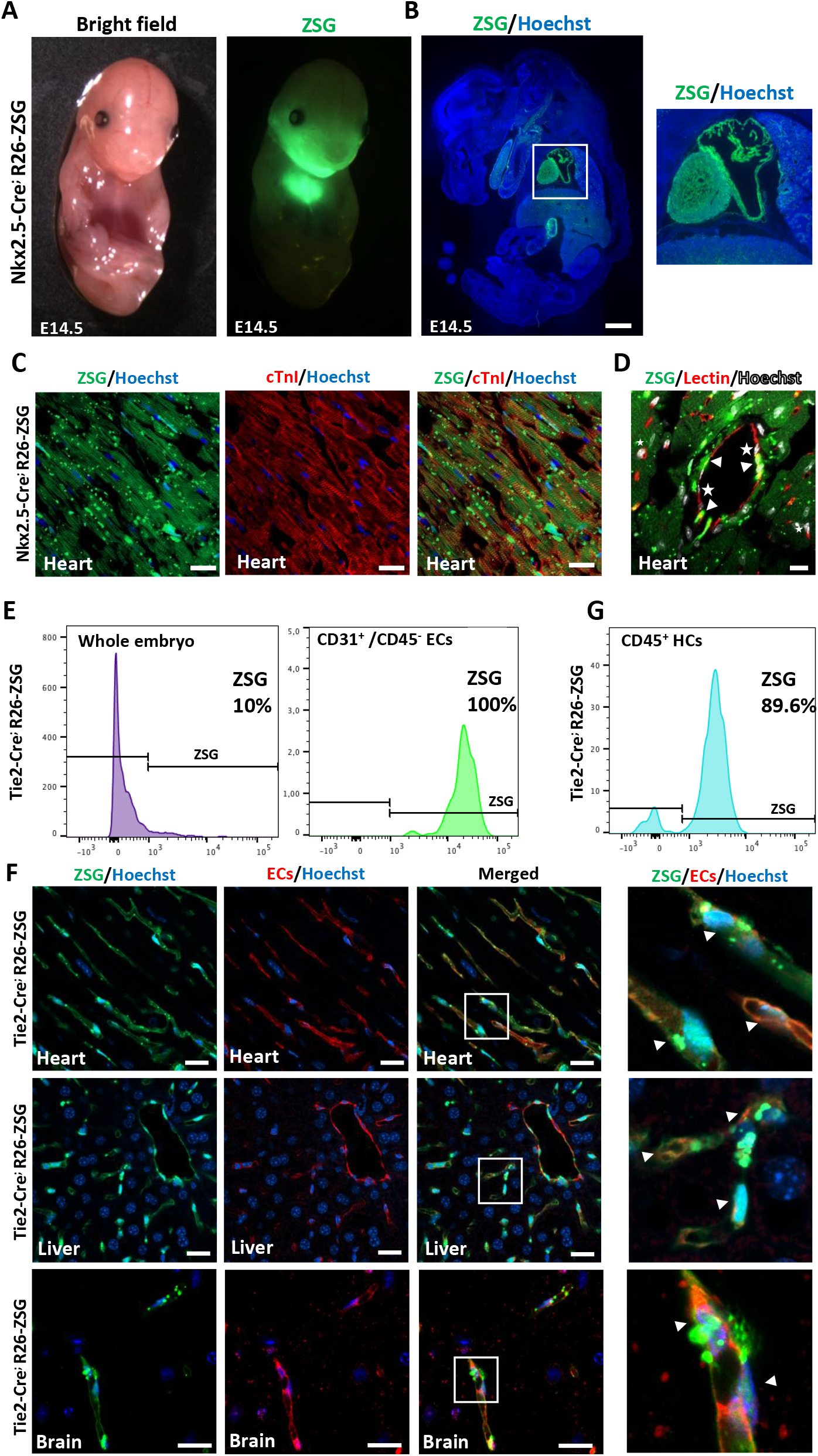
Characterization of Nkx2.5-Cre; R26-ZSG and Tie2-Cre; R26-ZSG reporter mice. (A-B) ZSG expression in a E14.5 Nkx2.5-Cre; R26-ZSG embryo: macroscopic view (A) and sagittal section (B). The insert shows a magnification of the heart region; scale bar 1mm. (C) Representative images of cTnI staining (in red) in the adult heart of a Nkx2.5-Cre; R26-ZSG mouse (in green). Nuclear Hoechst in blue. Scale bar 20μm. (D) BS-I Lectin staining (in red) in the adult heart of a Nkx2.5-Cre; R26-ZSG mouse (ZSG in green). Nuclear Hoechst in white. Scale bar 10μm. Arrowheads indicate ZSG^+^ ECs and stars ZSG^-^ ECs. (E) Representative histograms from flow cytometry analysis of ZSG^+^cells in the whole embryo (left) and in the ECs (CD31^+^; CD45^-^) (right) of Tie2-Cre; R26-ZSG embryos at E14.5. (F) Representative images of ECs immunofluorescence stainings (BS-I Lectin or CD31, in red) and ZSG expression (in green) in the adult organs of Tie2-Cre; R26-ZSG mice. The inserts (right) show a magnification of the selected areas (white squares). Scale bars 20μm. (G) Representative histogram from flow cytometry analysis of ZSG^+^ cells in the hematopoietic cell population (CD31^-/low^;CD45^+^) of whole blood of adult Tie2-Cre; R26-ZSG mice.

**Table S1:** Intraspecies blastocyst complementation experiments.

| Embryo donor strains | Cell line | N° microinjected embryos | N° of recipients | Developmental staged analyzed | N° of live embryos/mice (% of transferred embryos) | N° of chimera (% of live embryos/mice) | Nkx2.5-Cre; R26-DTA complemented (% of genotyped chimeras) | Tie2-Cre; R26-DTA complemented (% of genotyped chimeras) | Nkx2.5-Cre; Tie2-Cre; R26-DTA complemented (% of chimeras) | Nkx2.5-Cre; R26-DTA complemented (% of acardiac microinjected embryos) | Tie2-Cre; R26-DTA complemented (% of avascular microinjected embryos) | Nkx2.5-Cre; Tie2-Cre; R26-DTA (% of acardiac and avascular microinjected embryos) |
| --- | --- | --- | --- | --- | --- | --- | --- | --- | --- | --- | --- | --- |
| Nkx2.5-Cre x R26-DTA | AHFiPS 7 | 239 | 18 | E14.5 | 8 (3.3) | 8 (100) | 8 (100) | n.a. | n.a. | 8 (3.3) | n.a. | n.a. |
| Nkx2.5-Cre x R26-DTA | AHFiPS 9 | 81 | 6 | E14.5 | 5 (6.2) | 5 (100) | 5 (100) | n.a. | n.a. | 5 (6.2) | n.a. | n.a. |
| Nkx2.5-Cre x R26-DTA | R1 | 20 | 2 | E14.5 | 1 (5) | 1 (100) | 1 (100) | n.a. | n.a. | 1 (5) | n.a. | n.a. |
| Nkx2.5-Cre x R26-DTA | TOTAL | 340 | 26 | E14.5 | 14 (4.1) | 14 (100) | 14 (100) | n.a. | n.a. | 14 (4.1) | n.a. | n.a. |
| Tie2-Cre x R26-DTA | AHFiPS 7 | 290 | 18 | E14.5 | 44 (15.2) | 28 (63.6) | n.a. | 0 (0.0) | n.a. | n.a. | 0 (0.0) | n.a. |
| Tie2-Cre x R26-DTA | AHFiPS 9 | 92 | 6 | E14.5 | 15 (16.3) | 14 (93.3) | n.a. | 5 (35.7) | n.a. | n.a. | 5 (10.9) | n.a. |
| Tie2-Cre x R26-DTA | TOTAL | 382 | 24 | E14.5 | 59 (15.4) | 42 (71.2) | n.a. | 5 (11.9) | n.a. | n.a. | 5 (2.6) | n.a. |
| Nkx2.5-Cre; Tie2-Cre x R26-DTA | AHFiPS 9 | 326 | 18 | E14.5 | 14 (4.2) | 14 (100) | 12 (85.7) | n.a. | 2 (14.3) | 12 (7.4) | n.a. | 2 (1.2) |
| Nkx2.5-Cre; Tie2-Cre x R26-DTA | AHFiPS 7 | 110 | 6 | newborn/adult | 2 (1.8) | 2 (100) | 2 (100) | n.a. | 0 (0.0) | 2 (3.6) | n.a. | 0 (0.0) |
| Nkx2.5-Cre; Tie2-Cre x R26-DTA | AHFiPS 9 | 699 | 45 | newborn/adult | 11 (1.6) | 11 (100) | 5 (50)* | n.a. | 5 (50)* | 5 (1.4) | n.a. | 5 (1.4) |
| Nkx2.5-Cre; Tie2-Cre x R26-DTA | R1 | 395 | 16 | newborn/adult | 23 (5.8) | 23 (100) | 14 (70)* | n.a. | 6 (30)* | 14 (7) | n.a. | 6 (1.5) |
| Nkx2.5-Cre; Tie2-Cre x R26-DTA | TOTAL | 1204 | 67 | newborn/adult | 36 (3.0) | 36 (100) | 21 (65.6)* | n.a. | 11 (34.4)* | 21 (3.5) | n.a. | 11 (1.8) |

\* genotype of 1 chimera complemented with AHF9 non determined (ND); with R1 3 ND

**Table S2:**
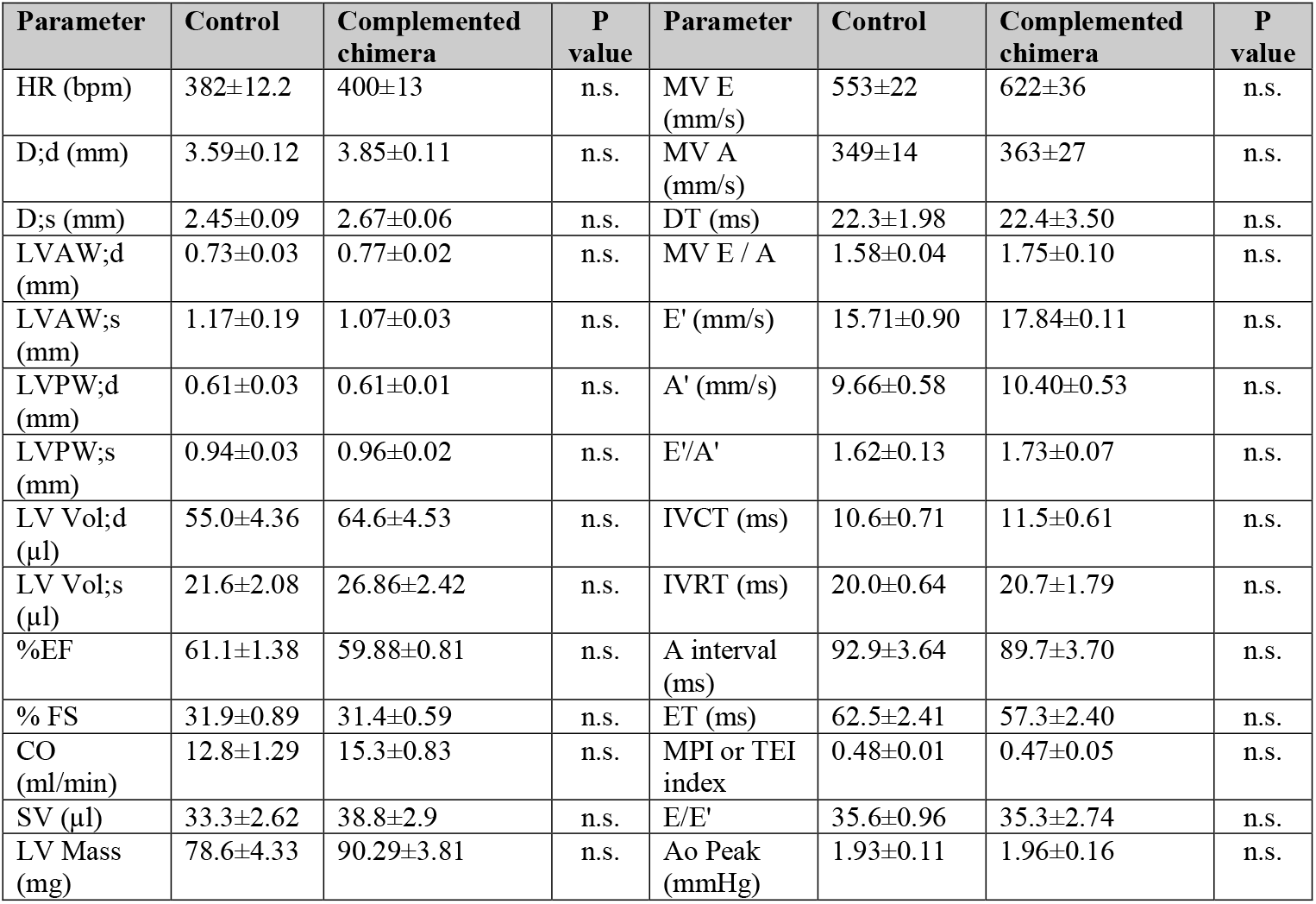
echocardiographic measurements in complemented adult chimeras.

**Table S3:** Interspecies blastocyst complementation experiments.

| Embryo donor strains | Cell line | N° microinjected embryos | N° of recipients | Developmental staged analyzed | N° of live embryos/ mice (% of transferred embryos) | N° of chimera (% of live embryos/mice) | Nkx2.5-Cre; R26-DTA complemented (% of genotyped chimeras) | Nkx2.5-Cre; Tie2-Cre; R26-DTA complemented (% of chimeras) | Nkx2.5-Cre; R26-DTA complemented (% of acardiac microinjected embryos) | Nkx2.5-Cre; Tie2-Cre; R26-DTA (% of acardiac and avascular microinjected embryos) |
| --- | --- | --- | --- | --- | --- | --- | --- | --- | --- | --- |
| Nkx2.5-Cre; Tie2-Cre x R26-DTA | ATCrSD13 | 116 | 11 | E10,5 | 15 (12.9) | 6 (40) | 0 (0) | 0 (0) | 0 (0) | 0 (0) |
| Nkx2.5-Cre; Tie2-Cre x R26-DTA | ATCrSD27 | 30 | 4 | E10,5 | 4 (13.3) | 2 (50) | 1 (50) | 0 (0) | 1 (6.6) | 0 (0) |
| Nkx2.5-Cre; Tie2-Cre x R26-DTA | ATCrSD28 | 50 | 4 | E10,5 | 2 (4) | 0 (0) | 0 (0) | 0 (0) | 0 (0) | 0 (0) |
| Nkx2.5-Cre; Tie2-Cre x R26-DTA | ATCrSD30 | 82 | 9 | E10,5 | 10 (12.2) | 6 (60) | 3 (50) | 0 (0) | 3 (7.3) | 0 (0) |
| Nkx2.5-Cre; Tie2-Cre x R26-DTA | TOTAL | 278 | 28 | E10,5 | 31 (11.2) | 14(45.2) | 4 (28.6) | 0 (0) | 4 (2.8) | 0 (0) |
| Nkx2.5-Cre; Tie2-Cre x R26-DTA | ATCrSD29 | 83 | 8 | E11,5 | 0 (0) | 0 (0) | 0 (0) | 0 (0) | 0 (0) | 0 (0) |
| Nkx2.5-Cre; Tie2-Cre x R26-DTA | TOTAL | 83 | 8 | E11,5 | 0 (0) | 0 (0) | 0 (0) | 0 (0) | 0 (0) | 0 (0) |
| Nkx2.5-Cre x R26-DTA | ATCrSD78 | 209 | 15 | E14,5 | 0 (0) | 0 (0) | 0 (0) | n.a. | 0 (0) | n.a. |
| Nkx2.5-Cre x R26-DTA | TOTAL | 209 | 15 | E14,5 | 0 (0) | 0 (0) | 0 (0) | n.a. | 0 (0) | n.a. |

**Table S4:** Antibodies list.

| **Primary Antibody** | **Source** | **Catalog #** | **Dilution** |
| --- | --- | --- | --- |
| anti-mouse CD31 | BD Pharmigen | 557355 | 1:50 |
| BS-I-lectin-Biotin | Sigma-Aldrich | L3759 | 1:100 |
| anti-Troponin I | Santa Cruz | Sc-133117 | 1:100 |
| anti-Nkx2.5 | Santa Cruz | Sc-8697 | 1:150 |
| anti-GFP | Invitrogen | A11122 | 1:200 |
| **Secondary Antibody** | **Source** | **Catalog #** | **Dilution** |
| Goat anti-rat Alexa 594 | Invitrogen | A11007 | 1:500/1:200 |
| Goat anti-mouse Alexa 594 | Invitrogen | A11032 | 1:500/1:200 |
| Donkey anti-goat 594 | Invitrogen | A11058 | 1:1000 |
| Goat anti-rabbit 488 | Invitrogen | A11008 | 1:1000 |
| Streptavidin-HRP | PerkinElmer | NEL750001EA | 1:75 |
| Streptavidin-Cy3 | Sigma-Aldrich | GEPA43001 | 1:200 |
| Rabbit anti-rat IgG, H and L-Biotin | Abcam | ab6733 | 1:200 |
| **FACS Antibody** | **Source** | **Catalog #** | **Dilution** |
| anti-CD31APC | BioLegend | 102420 | 1:100 |
| anti-CD45-PE | BD Bioscience | 553081 | 1:100 |
| rat IgG2a APC | BioLegend | 400512 | 1:100 |
| rat IgG2b PE | BD Bioscience | 553989 | 1:100 |

**Table S5:**
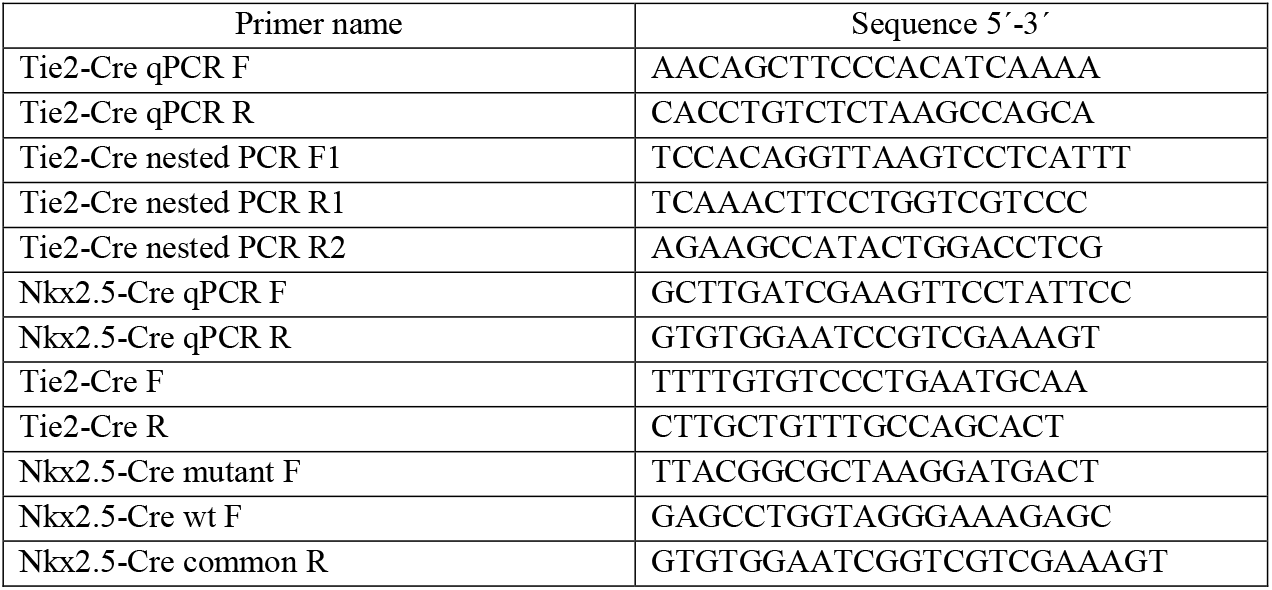
Primers list.

## Notes

### Competing Interest Statement

The authors have declared no competing interest.

